# MATING FREQUENCY MEDIATES PERSONALITY EXPRESSION IN FACULTATIVELY POLYANDROUS MITES

**DOI:** 10.1101/2024.12.23.630184

**Authors:** Peter Schausberger, Shogo Usugi, Chenhao Wang, Norihide Hinomoto

## Abstract

Animal personalities are characterized by within-individual consistency linked to among-individual variability. Personality expression may change during ontogeny and is often dependent on major life history transitions and events such as mating and the onset of reproduction. While the influence of female personalities on sexual selection and mating has long been known, the influence of the females’ mates on female personality expression after mating is poorly understood. Here we hypothesized that in facultatively polyandrous animals multiple mating increases the females’ assets, also called the residual reproductive value (RRV), due to direct and/or indirect benefits. Based on the predictions of the asset protection principle, higher RRV should promote behaviors that reduce the risk of fitness loss and hence mediate behavioral repeatability displayed in groups. We tested our hypothesis in the group-living, plant-inhabiting predatory mite *Phytoseiulus persimilis*. To this end, predatory mite females were presented with either a single mate or two mates in sequence and their post-mating repeatability in activity and sociability was evaluated in groups that were composed of females of the same and mixed mating types (monandrous and polyandrous). Mating frequency had little effects on activity patterns but pronounced effects on sociability traits. Polyandrous females were on average more sociable as well as more repeatable in sociability than monandrous females. These behavioral shifts reflect greater risk aversion and strategies to mitigate inter-individual conflicts within groups to enhance asset protection. Our study suggests that the mating frequency can critically influence female personality expression after mating in facultatively polyandrous animals and highlights the importance of considering mate-related variables in animal personality research.

## Introduction

Animal personalities are characterized by consistent within-individual behaviors that consistently differ among individuals across time and contexts^1,2^. Animal personalities are not bound to certain taxonomic groups but are observed across animals, from anemones, arachnids, and insects to birds, fish and mammals^3,4^. Typical pathways mediating personality expression are genetic determination, transgenerational information transfer, and personal experiences. In many animals, personalities are not fixed throughout life but change over the course of ontogeny and/or are contingent upon major life history transitions or events^5^, such as when being released from parental care, when becoming adult, when mating and starting to reproduce or when ceasing reproduction^6,7,8^.

The influence of personality expressed before or during mating on sexual selection and mating are relatively well researched^9,10^, but the influence of mating-associated events on female personality expression after mating remains elusive. Mating-related circumstances such as mate search and choice, the mating event itself, mating frequency and the types of mates are some of the most critical events for any sexually reproducing animal, for being decisive determinants of reproductive success and fitness. However, the proximate and ultimate aspects of mating-mediated adjustments in personality expression are poorly understood. This is also true for the role of the quality and/or quantity of male mates on female personality expression, yet the few studies that are available suggest a potentially larger influence than anticipated. For example, Monestier & Bell (2020)^11^ showed that both mating and mere presence of a courting male critically change mean expression and repeatability of boldness and sociability of stickleback females, *Gasterosteus aculeatus*, as compared to no mating experience. Schausberger & Nguyen (2024)^12^ observed that the early-life social experience of male mates critically influences repeatability of activity and sociability of predatory mite females after mating. In polyandrous animals, initial mate choice and first mating experience are often decisive for subsequent choice and re-mating willingness^13^. Females typically become more selective with mating experience, which in turn should affect their personality expression. The influence of mating frequency on female personality expression is unknown for any animal.

Powerful ultimate explanations for mating-related adjustments of female personalities are provided by the predictions of the asset protection principle^14^. The asset protection principle postulates that individuals with many assets, i.e. high residual reproductive value (RRV), should avoid risks to allow harvesting these assets, while individuals with few assets, i.e. low RRV, should take more risks because they have less to lose^14^. Accordingly, mating-related experiences should influence mean behavioral trait expression as well as personality traits such as boldness (risk-taking behavior), because of the trade-off between current and future reproduction^15^. While the asset protection principle has been originally developed^14^ and evaluated^16,17^ for predation risk taking, it is applicable to any behavior that is fitness relevant and bears some kind of risk of fitness loss/decrease^15^, including the canonical personality traits aggressiveness and sociability, especially in group-living animals.

Here we tested the hypotheses that in facultatively polyandrous animals, which is a widespread mating system among animals^13,18-20^, multiple mating increases the assets (RRV), compared to single mating, and should thus lead to more risk-averse behaviors (in a broad sense: risk of fitness loss). Multiple mating can increase the assets through direct (material) and/or indirect (genetic) benefits^13,21^. Multiple mating should diversify the females’ behaviors, because each female experiences a different set of two mates with presumably different qualities, compared to single mating, which could in turn result in lower within-individual variability relative to among-individual variability, together increasing behavioral repeatability. Higher assets (higher RRV) should be linked to increased behavioral repeatability if the assets cannot be increased anymore (no search or choice of additional mates needed, no mate competition, diversity among females is promoted because different sets of two mates provide more complex information to females and allow much more variability than single mates). Increased assets should result in more consistent within-individual behaviors, relative to among-individual variability, if within-individual consistency coupled to among-individual variability and asset value are linked by positive feedback loops^22-24^.

We tested our hypotheses in plant-inhabiting predatory mites *Phytoseiulus persimilis*. Theses mites live in groups, which is brought about by the patchy distribution of their prey (spider mites) and mutual attraction^25-28^. *P. persimilis* females are facultatively polyandrous, that is, they mate once or twice. Monandrous and polyandrous females produce similar numbers of eggs, but the second mate provides indirect benefits, due to offspring sired by two males^29^. Having an additional mate increases the assets of *P. persimilis* females because their offspring become genotypically and phenotypically more diverse^29^, andd thus constitute as a whole a higher RRV for their mothers.

## Materials and methods

### Predatory mite origin and rearing

Predatory mites *Phytoseiulus persimilis* used in experiments derived from a laboratory-reared population founded with specimens collected on eggplant, *Solanum melongena*, in Sicily^12^. In the laboratory, the predatory mites were reared in heaps of detached leaves of common bean plants, *Phaseolus vulgaris*, infested with two-spotted spider mites *Tetranychus urticae*, inside small plastic boxes (11.7 x 16.5 x 5.6 cm). The small plastic boxes were fixed inside large plastic boxes (23.9 x 17.6 x 9.1 cm), containing a shallow layer of soapy water, and closed on top by a lid with a mesh-covered rectangular opening (8 x 10 cm) for ventilation. The rearing boxes were stored in an air-conditioned room at 25±1 °C, 60 to 80% RH and 16:8 h L:D. The spider mite population was founded by specimens collected from chrysanthemum plants, *Chrysanthemum morifolium*, in Nara, Japan and reared on whole common bean plants at room temperature under daylight fluorescent lamps (16:8 h L:D). Spider mite-infested bean leaves were clipped off the plants and added to the small boxes used for rearing the predatory mites twice per week.

### Pre-experimental treatments

To generate predatory mite females used in experiments, gravid females were randomly withdrawn from the rearing and placed in groups of 50 on detached spider mite-infested bean leaf arenas. Each arena consisted of a primary leaf (∽ 8 x 8 cm) placed upside down on moist filter paper on top of a circular foam pad resting inside a Petri dish (9 cm Ø) half-filled with water. Moist tissue paper was wrapped around the edges of the leaf arena. Predatory mite eggs <24h old were collected and transferred in groups of 50 to fresh large leaf arenas, infested with spider mites, for development; leaf arenas were monitored once per day until the predatory mites had reached the deutonymphal stage. Female deutonymphs were removed and singly placed on circular bean leaf discs (1.2 to 1.5 cm Ø), harboring mixed spider mite stages, resting upside down on water agar columns (1.2 to 1.5 cm Ø, 1.1 cm high) inside closed acrylic cages half-filled with water. Each acrylic cage consisted of a closed Petri dish (5.0 cm Ø, height 1.5 cm) with a mesh-covered (mesh size 0.05 mm) ventilation opening (1.3 cm Ø) in the lid (SPL Life Sciences Co. Ltd., S-Korea). As soon as the females were adult, one adult male, randomly taken from another large leaf arena than the female, was added to the leaf disc and left there for 3 h. Mating typically commences within 30 min after pairs are together on a leaf and lasts on average around 120 to 180 min^29-31^. After 5 to 6 h, the male was removed while the female was left on the disc. After another two days, half of the females received a second male for mating for 5 to 6 h (to represent polyandrous females in the experiment), while the other half of females was left without a second male (to represent monandrous females in the experiment). Sample sizes at the start of the behavioral assays were 37 monandrous females and 50 polyandrous females. Eggs were counted and removed from the discs in 1 to 2 d intervals. All experimental units were stored in an air-conditioned room at 25 ± 1 °C, 60 to 80% RH and 16:8 h L:D.

### Behavioral assays

Each monandrous and polyandrous female was subjected to two group assays, dubbed pure and mixed, using rectangular acrylic arenas (2.0 x 1.1 cm). Each arena was bordered by 1 to 2 mm high walls, built of water-soaked cotton pads, and pre-loaded with ∽60 eggs of two-spotted spider mites, spread equally on the surface of the arena before conducting the assay. In pure group assays, four females with the same mating type were grouped together in each arena (either four monandrous or four polyandrous females). In mixed group assays, two monandrous and two polyandrous females were grouped together in each arena. Before assays, each predatory mite female was uniquely colored by a small watercolor dot on her dorsal shield to make her distinguishable during the assays. To start a group assay, four differently colored females were transferred from their leaf discs in the arena and allowed to acclimatize for 5 min. After 5 min, the behavior of the females was videotaped for 4 min using either a USB digital microscope (Jiusion HD 2MP, Shenzhen Jiu Sheng Electronic Commerce, Shenzhen, China) or a color CMOS camera (Wraycam NOA2000, Wraymer, Osaka, Japan) attached to a stereo microscope (Olympus SZ61TRC-C, Tokyo, Japan). After the group assays, the females were immediately returned to the leaf disc they came from. Mixed group assays took place two days after the pure group assays (i.e., females had a 2 d rest in between the pure and mixed group assays).

### Video-tracking and statistical analysis

Videotaped behaviors (15 frames/s in each video) were automatically analyzed using AnimalTA, version 3.2.1^32^. AnimalTA allows tracking the movement and inter-individual interactions separately for each individual of a group. The trajectories of each individual target obtained by video-tracking were used for analysis of the proportion of time moving (moving threshold was 0.2 mm/s), running speed when moving, inter-individual distance, number of immediate neighbors (i.e., within a radius of 2 mm from the target, which is the touching distance when both interactants longitudinally stretch their first pair of legs^33^), and the proportion of time spent with at least 1 neighbor.

All statistical analyses were conducted using IBM SPSS Statistics v. 29.0.1^34^. Separate generalized estimating equations (GEEs; linear) were used to analyze the effects of female mating frequency (mono- or polyandrous) and type of group assay (pure or mixed) on the mean proportion of time spent moving, running speed, inter-individual distance, number of neighbors and proportion of time with at least 1 neighbor. A generalized linear model (GLM; linear) was used to examine the effect of female mating frequency on the number of eggs produced on their leaf discs over 5 d. Personality expression by monandrous and polyandrous females, i.e. behavioral repeatability between group assays, in the proportion of time spent moving, running speed, inter-individual distance, number of neighbors and proportion of time with at least one neighbor was evaluated by intraclass correlation coefficients (ICC; two-way random, consistency, average measure^35^). Within- and among-individual variances were checked to pinpoint the causes of different ICCs induced by female mating frequency (mono- or polyandrous). SuperPlotsOfData^36^ was used to create figures 1 and 2.

**Figure 1.**
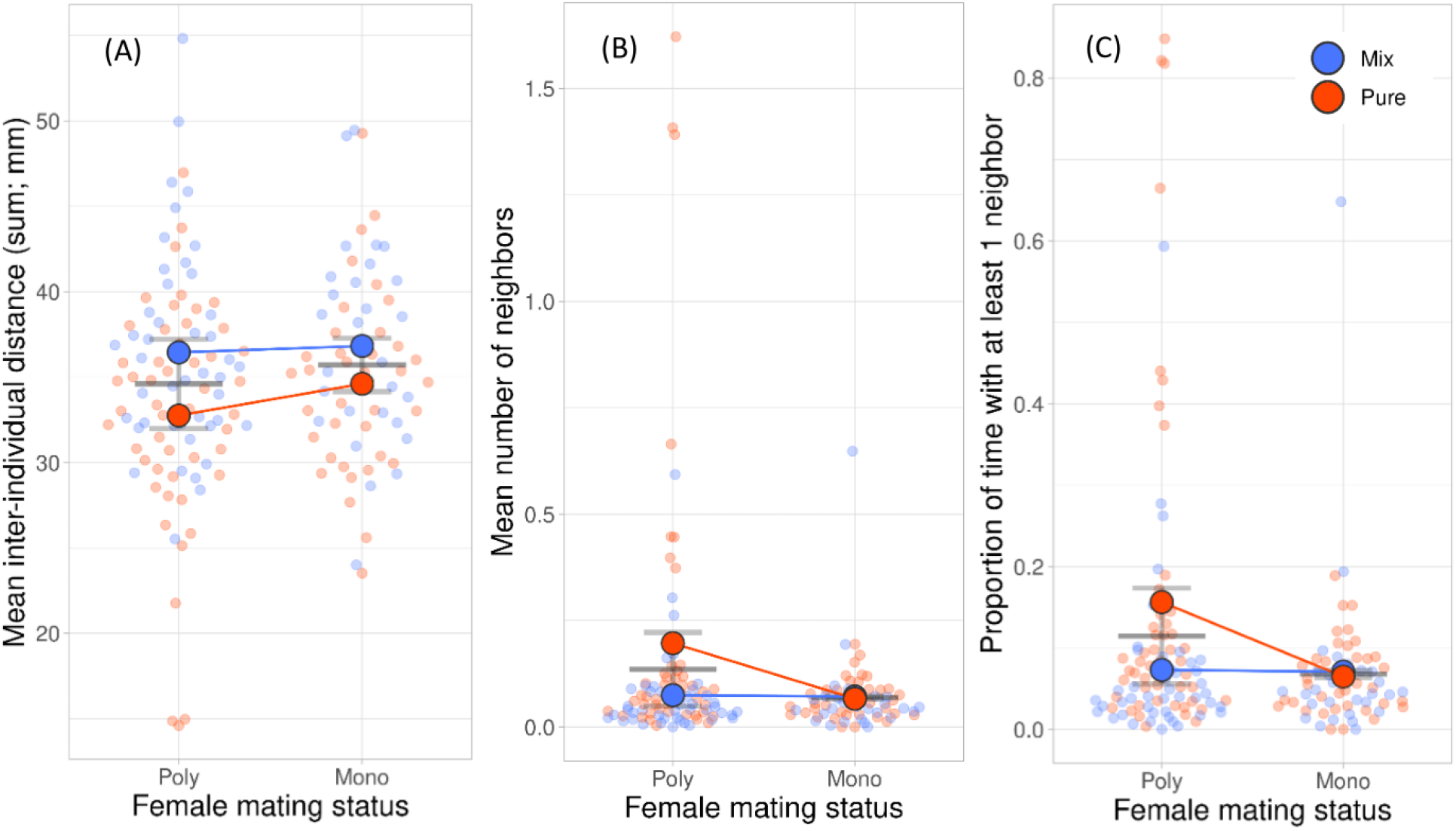
Mean (± SE) inter-individual distance (A), mean number of neighbors (B; within a radius of 2 mm of the target) and mean proportion of time with at least one neighbor (C) of *Phytoseiulus persimilis* females, which had mated once (mono) or twice (poly), sequentially observed in groups of four females with the same (pure) and different (mix) mating frequency. Pale dots are the individual data.

**Figure 2.**
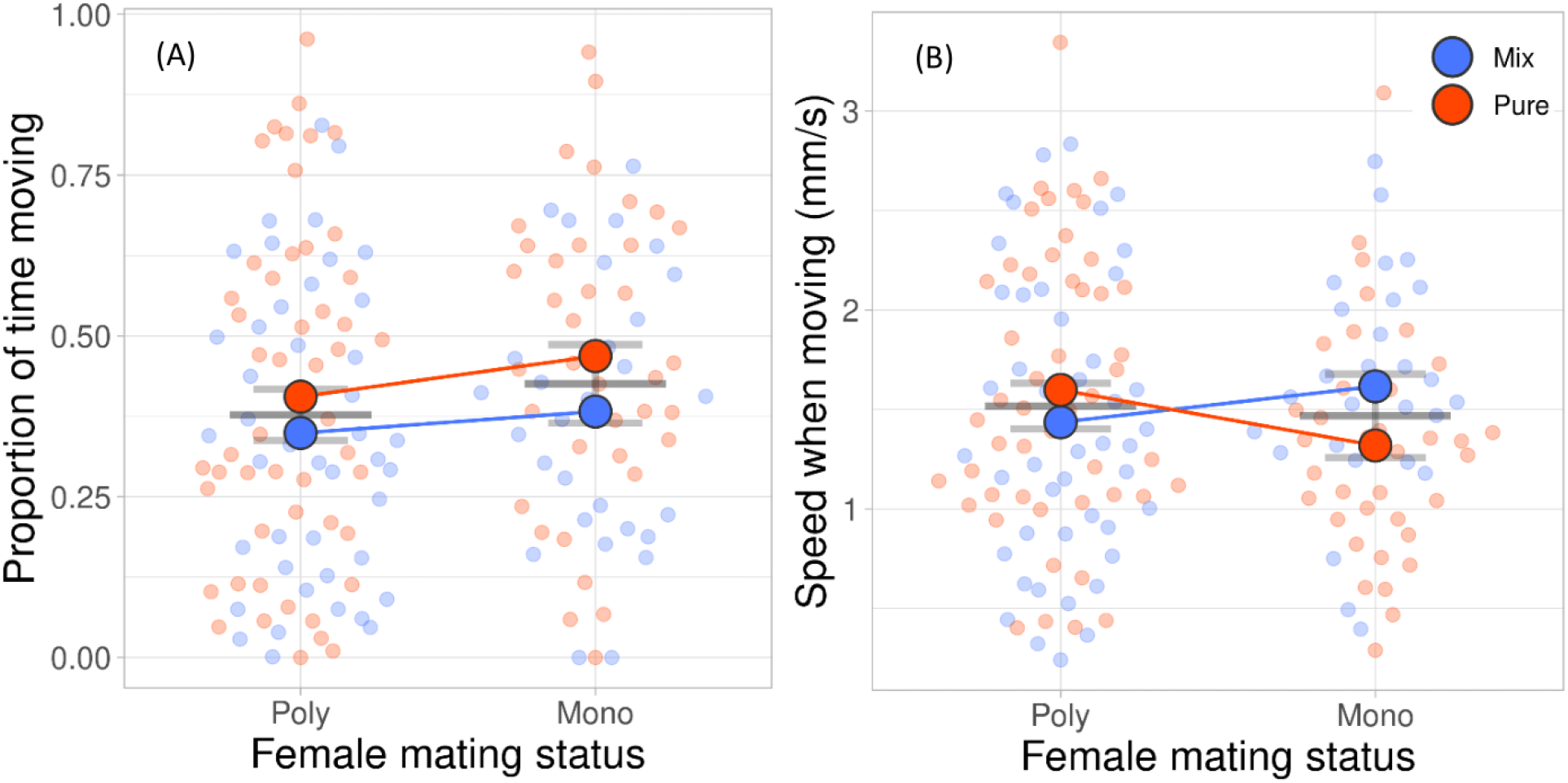
Mean (± SE) proportion of time moving (A) and mean running speed when moving (B) of *Phytoseiulus persimilis* females, which had mated once (mono) or twice (poly), sequentially observed in groups of four females with the same (pure) and different (mix) mating frequency. Pale dots are the individual data.

## Results

Type of group assay (pure or mixed group) had a significant main effect on the mean inter-individual distance, number of immediate neighbors (within a radius of 2 mm of the center of the target), proportion of time with at least one neighbor and proportion of time moving of *Phytoseiulus persimilis* females (Table 1, Fig 1ABC, 2A). Females were closer together (Fig 1A), had more neighbors (Fig 1B), spent more time with at least one neighbor (Fig 1C) and moved more (Fig 2A) in pure than mixed groups. However, having more neighbors and spending more time with at least one neighbor in pure than mixed groups was only the case in polyandrous but not monandrous females (Fig 1BC). The number of eggs produced on the leaf discs did not differ between monandrous (11.79 ± 0.80 SE) and polyandrous (12.50 ± 0.66 SE) females (GLM: Wald chi-square = 0.460, *P* = 0.498). Polyandrous females ran faster in pure than mixed groups whereas the opposite was true for monandrous females (Fig 2C). Polyandrous but not monandrous females were repeatable in inter-individual distance, number of immediate neighbors and time spent with at least one neighbor (Table 2). Neither type of female was repeatable in the proportion of time moving and running speed (Table 2). Scrutiny of the among- and within-individual variances of the sociability traits revealed that the higher repeatability of polyandrous than monandrous females was due to: (i) for inter-individual distance, a combination of higher among-individual variability and higher within-individual consistency and smaller within- than among-variability in polyandrous but not monandrous females, (ii) for the number of neighbors and the proportion of time with at least 1 neighbor, smaller within- than among-variability in polyandrous females, whereas the within- and among-variabilities were similar in monandrous females. The variances in sociability traits had different levels in polyandrous and monandrous females, with higher levels in polyandrous females. Both variances were extremely small and similar in monandrous females, suggesting that all females behaved very similar; polyandrous females diversified more among each other, but behaved, in relative comparison of among and within-variability, more consistent within individuals.

**Table 1.** Results of generalized estimating equations (GEE) of the effects of the mating frequency (once or twice) of *Phytoseiulus persimilis* females and the type of group assay (pure or mixed). Each female was sequentially observed in a group of four females with the same (pure; all either monandrous or polyandrous) and different (mixed; two monandrous and two polyandrous) mating status.

| Dependent variable | Independent variables | N | Wald<br>Chi-square | df | P-value |
| --- | --- | --- | --- | --- | --- |
| Inter-individual distance | (Intercept) | 161 | 5222.209 | 1 | <.001 |
|  | Mating frequency |  | 1.423 | 1 | 0.233 |
|  | **Type of assay** |  | **10.663** | **1** | **0.001** |
|  | Mating frequency*type of test |  | 0.698 | 1 | 0.404 |
| Number of neighbors | (Intercept) | 161 | 41.577 | 1 | <.001 |
|  | **Mating frequency** |  | **4.454** | **1** | **0.035** |
|  | **Type of assay** |  | **5.637** | **1** | **0.018** |
|  | **Mating frequency\*type of assay** |  | **6.805** | **1** | **0.009** |
| Time with $\geq 1$ neighbor | (Intercept) | 161 | 64.132 | 1 | <.001 |
|  | **Mating frequency** |  | **4.152** | **1** | **0.042** |
|  | **Type of assay** |  | **4.799** | **1** | **0.028** |
|  | **Mating frequency\*type of assay** |  | **6.584** | **1** | **0.010** |
| Time moving | (Intercept) | 161 | 463.104 | 1 | <.001 |
|  | Mating frequency |  | 1.657 | 1 | 0.198 |
|  | **Type of assay** |  | **4.062** | **1** | **0.044** |
|  | Mating frequency*type of assay |  | 0.200 | 1 | 0.655 |
| Running speed | (Intercept) | 157 | 850.683 | 1 | <.001 |
|  | Mating frequency |  | 0.208 | 1 | 0.648 |
|  | Type of assay |  | 0.476 | 1 | 0.490 |
|  | **Mating frequency\*type of assay** |  | **5.362** | **1** | **0.021** |

Significant effects ( $P < 0.05$ ) are highlighted in bold

**Table 2.** Intraclass correlation coefficients (ICC, average measure; repeatability) of behavioral traits of *Phytoseiulus persimilis* females as affected by the mating frequency (mono- or polyandrous). Each female was sequentially observed in a group of four females with the same (pure; all either monandrous or polyandrous) and mixed (two monandrous and two polyandrous) mating status.

| Personality trait | Mating frequency | N | ICC | CI (95%) | P-value |
| --- | --- | --- | --- | --- | --- |
| Inter-individual distance | Mono | 28 | -0.272 | -1.750; 0.411 | 0.732 |
|  | **Poly** | **44** | **0.497** | **0.078; 0.726** | **0.013** |
| Number of neighbors | Mono | 28 | -0.098 | -1.374; 0.492 | 0.595 |
|  | **Poly** | **44** | **0.493** | **0.070; 0.723** | **0.014** |
| Time with $\geq 1$ neighbor | Mono | 28 | -0.080 | -1.334; 0.500 | 0.579 |
|  | **Poly** | **44** | **0.578** | **0.227; 0.770** | **0.003** |
| Time moving | Mono | 28 | -0.039 | -1.245; 0.519 | 0.539 |
|  | Poly | 44 | 0.233 | -0.407; 0.581 | 0.195 |
| Running speed | Mono | 26 | 0.379 | -0.384; 0.722 | 0.120 |
|  | Poly | 42 | -0.306 | -1.430; 0.298 | 0.802 |

Significant effects ( $P < 0.05$ ) are highlighted in bold

## Discussion

Our study documents that the mating frequency mediates personality expression in facultatively polyandrous predatory mites *Phytoseiulus persimilis*. Polyandrous but not monandrous females were repeatable in sociability traits such as inter-individual distance, number of immediate neighbors and proportion of time spent with at least one neighbor. High repeatability in sociability traits of polyandrous females was primarily due to lower within-individual variances than among-individual variances. In monandrous females and depending on the trait, within and among-variances were similar or among-variance was higher than within-individual variances, compromising repeatability. Polyandrous females were not only more repeatable but on average also more sociable than monandrous females. The type of group composition (purely monandrous/polyandrous or mixed) influenced the mean behavioral trait expressions in activity and sociability traits, but more so in polyandrous than monandrous females.

Mating frequency was a previously unknown factor mediating the behavioral repeatability of females following mating. Other mating-related factors were documented for sticklebacks^11^ and the very same animals of this study, predatory mites *Phytoseiulus persimilis*^12^. Sticklebacks became, on average, less sociable and less risk-taking after mating or mere courtship by males; also, repeatability of sociability and boldness before and after the mating opportunity were lower than in the unmated control group^11^. Female sticklebacks compete for placing their eggs inside the nests built by males, which may explain why they became less sociable and more competitive after mating. Our study animals, the group-living *P. persimilis*, became both on average more sociable and more repeatable in sociability following the second mating. Such behavioral shifts should enhance fitness in group-living species. Reproducing *P. persimilis* females benefit from collective foraging and spider mite patch exploitation, with these benefits being higher than the costs of competition. Group-living is the norm in *P. persimilis*, only during dispersal and low prey availability, or when the previously ample prey supply deteriorates, the groups dissolve.

From a proximate perspective, the stickleback study by Monestier & Bell (2020)^11^ suggests that the mere social experience of being courted by a male can alter the repeatability of female behavior, with the social and sexual (mating) experiences exerting the same effects in the sticklebacks. Our experimental procedure presented the females one (for the monandrous treatment) or two males (for the polyandrous treatment), with the duration of each male presentation allowing only for one complete mating, typically lasting 2 to 3 h^29,35^. All females mated once, as indicated by egg production after the first mating. All females of the polyandry treatment experienced the presence of a second mating partner and we assume that most of them, if not all, did mate twice. Typically, around 90% of *P. persimilis* females do mate upon presentation of a second mate, with around 70% of these producing offspring with mixed paternities^29^.

Ultimate explanations for why multiple mating produced more repeatable females (more variable among individuals but, in relative comparison, more consistent within individuals) can be inferred from mate competition and asset (RRV) protection^14^. Mate competition is over for polyandrous females, whereas monandrous females still compete for additional mates. This also explains why polyandrous females were on average more sociable in groups consisting of just polyandrous females but not in mixed groups, consisting of both monandrous and polyandrous females. Polyandrous females are not engaging anymore in mate competition with other females and thus have lower needs to flexibly adjust their behavior to the behavior of others. Due to their high RRV, polyandrous females should be more risk averse than monandrous females and behave in a way that limits the risk of fitness loss. They should be more protective of their offspring than monandrous females and optimize their realized niches, including the interactions with conspecifics. By predictably signaling their state to other co-habiting females, they can avoid the costs of mate competition; by signaling their state to cohabiting males, they can avoid the costs of mating attempts that would not result in any direct or indirect benefits. Higher variability among polyandrous than monandrous females was likely brought about by the influence of different sets of two mates, providing highly variable information to females. Increased among-individual variability, reflecting individualized female behaviors, should reduce intra-group conflicts. Similar to the large difference in repeatability of monandrous and polyandrous females in this study, Schausberger & Nguyen (2024)^12^ observed strongly differing repeatabilities in females mediated by the early-life experiences of their mating partners. Male mates that had been socially isolated early in life strongly increased the repeatability in activity of females that had been grouped early in life^12^.

In conclusion, our study underscores the critical importance of the females’ mates on female personality expression after mating. Together with Monestier & Bell (2020)^11^ and Schausberger & Nguyen (2024)^12^, our study demonstrates the sensitivity of adult female personality formation to mate-related circumstances, such as the quantity and phenotypes of mates, mediating mean trait expression as well as behavioral repeatability after mating. Our study suggests that mating frequency and other mating-related aspects deserve more attention in animal personality studies, for being potential drivers of behavioral repeatability in sexually reproducing animals, with significant consequent ecological and evolutionary implications.

## Author contributions

PS and NH conceived the study idea, developed the experimental design. acquired funding, provided resources and supervised the project. PS, SU, and CW conducted the study. PS analyzed the data and wrote the first draft of the manuscript. PS, SU, CW and NH contributed to revision, read and approved submission of the manuscript.

## Acknowledgements

We thank the University of Kyoto, Graduate School/Faculty of Agriculture for funding the stay of PS as an Invited Professor at the Laboratory of Ecological Information. This work was also financially supported by the Austrian Science Fund (FWF; P33787-B to PS).

